# sMEK1 promotes crosstalk between IRE1 and Akt signalling pathways: Evidence for a novel IRE1/sMEK1/Akt complex

**DOI:** 10.1101/2021.07.09.451832

**Authors:** Ozaira Qadri, Samirul Bashir, Mariam Banday, Debnath Pal, Khalid Majid Fazili

## Abstract

The Unfolded Protein Response (UPR) is a dynamic cellular pathway that helps maintain proteostasis during endoplasmic reticulum (ER) stress. One of the key UPR sensors is IRE1, which plays a central role in managing ER stress and interacts with other cellular pathways to regulate cell homeostasis. The Akt signalling pathway, on the other hand, is a crucial survival pathway involved in diverse cellular functions like growth, proliferation, glucose metabolism, and survival. This study explores the interplay between these two important cell signalling pathways. Specifically, our study revealed that IRE1 negatively regulates Akt through the protein phosphatase sMEK1. We identified sMEK1 and Akt as novel interacting partners of IRE1, which together form a ternary complex that helps coordinate the IRE1 and Akt signalling networks. The IRE1/sMEK1/Akt ternary complex results in the dephosphorylation of Akt by sMEK1 in the presence of activated IRE1. Together, this study sheds light on the molecular mechanism underlying the UPR/Akt link and provides valuable insights into the overall impact of their interaction.

## 1. Introduction

The Unfolded Protein Response (UPR) is a cellular stress response pathway that is activated in response to the accumulation of misfolded or unfolded proteins in the endoplasmic reticulum (ER). This pathway helps to restore ER homeostasis by reducing the load of protein synthesis and enhancing the clearance of misfolded proteins *(Hetz et al., 2011).* In higher eukaryotes, UPR is initiated by three main signalling proteins; Inositol requiring enzyme 1 (IRE1), double-stranded RNA dependent Protein kinase-like ER kinase (PERK) and Activating transcription factor 6 (ATF6) *(Schröder and Kaufman, 2005)*. Among these, IRE1 is the most evolutionary conserved and extensively studied arm of the UPR. IRE1 is a bifunctional enzyme with both kinase and endoribonuclease activity. When activated, IRE1 acts as an endoribonuclease, cleaving an mRNA encoding the transcription factor XBP1, which is then spliced to produce a functional protein that translocates to the nucleus and activates the expression of UPR genes, including chaperones, folding enzymes, and degradation factors *(Walter and Ron, 2011)*. IRE1 also degrades ER-targeted mRNAs in a process termed regulated IRE1-dependent mRNA decay (RIDD) *(Hollien and Weissman, 2006; Hollien et al., 2009)*. Besides behaving as a linear signalling network, IRE1 acts as a platform for docking several proteins that regulate IRE1 and bridge it to other cellular pathways. All these aspects of IRE1 make it a central molecule of the UPR signalling pathway that has a role in cell fate determination as well as in various diseases *(Bashir et al., 2021; Jager et al., 2012; Urra et al., 2020)*.

Several studies have put forth that UPR instead of being a unidirectional pathway, communicates with a number of other pathways and sends signals multi-directionally *(Jager et al., 2012)*. One of the key players is IRE1 which acts as the protein docking platform and orchestrates with other pathways to relay the signal. This concept is known as ‘UPRosome,’ where IRE1 is thought to act centrally and interact with a multitude of proteins that can either activate/inactivate or connect it with other signalling networks (*Hetz and Glimcher, 2009*; *Urano et al., 2000)*. The activating interactions of IRE1 include; Bcl-2 family members *(Hetz et al., 2006; Rodriguez et al., 2012*), Protein kinase C substrate 80K-H (PRKCSH) *(Shin et al., 2019)*, HSP72 *(Gupta et al., 2010),* HSP47 *(Sepulveda et al., 2018)*, HSP90 *(Marcu et al., 2002)*, Non-muscle myosin-IIb (NMIIB) *(He et al., 2012)* and Filamin *(Urra et al., 2018)*. All these proteins help in amplifying the IRE1 signal during ER stress. Other proteins like Fortilin *(Pinkaew et al., 2017)*, Bax inhibitor-1 (*Lisbona et al., 2009*), and Bid (*Bashir et al., 2023*) interact with IRE1 to inhibit its activity, while interaction with Nck, TNFR-associated factor 2 (TRAF2) *(Luo et al., 2008; Nguyen et al., 2004),* and RACK1 (*Qiu et al., 2010*) form the scaffolding interactions. Besides these physical interactions, IRE1 communicates with other cellular pathways, including the Akt/mTOR signalling pathway (*Ozcan et al. 2004, Qin et al., 2010, Wouters et al., 2008*).

Akt is a major survival factor promoting cell proliferation and whose dysregulation has been frequently detected in many types of cancer *(Liu et al., 2009).* Akt/PKB (Protein Kinase B) members are the major kinases that act downstream of PI3K. These are involved in various cellular functions, including growth, proliferation, glucose metabolism, invasion, metastasis, angiogenesis and survival (*Manning and Toker, 2011*). In this way, both Akt and UPR pathways are important for normal cellular function and survival, and any disruption in the two pathways can be the leading cause of various diseases, including cancer. Thus, studying the relay of signal and mechanistic crosstalk between these two pathways becomes essential in advancing our understanding of disease pathogenesis and potential therapeutic strategies.

Here we report that IRE1 and Akt share functional crosstalk through serine/threonine phosphatase, sMEK1. sMEK1 is a tumour suppressor protein also known as protein phosphatase 4 regulatory subunit 3 (PP4R3), a member of the PP2A subfamily, which has a conserved serine/threonine phosphatase domain *(Chen et al., 2008; Chowdhury et al., 2008; Nakada et al., 2008)*. sMEK1 acts as a novel pro-apoptotic protein while decreasing the expression of the AKT/mTOR pathway *(Byun et al., 2012; Kim et al., 2014; Kim et al., 2015)*. We found that IRE1 negatively regulates Akt phosphorylation through sMEK1. Our results revealed that both Akt and sMEK1 assemble in a complex with IRE1 which makes the basis for Akt regulation. Therefore, our results not only validate the connection between IRE1 and Akt pathways but also give a throughput understanding of the mechanism involved in the process.

## 2. Results

### 2.1 IRE1 interacts with a PP2A phosphatase sMEK1

The IRE1 protein is a key player in coordinating multiple signalling pathways within the cell. One of its fascinating functions is acting as a central docking platform for proteins to assemble and determine various cellular outputs. In order to investigate this, we conducted a mass spectrometry analysis to identify interacting partners of IRE1. Interestingly, our analysis revealed an enrichment of the sMEK1 protein (Supplementary Table S1). sMEK1 is known to be a critical protein in regulating numerous cellular processes, including growth, survival and metabolism *(Chen et al., 2008; Chowdhury et al., 2008; Nakada et al., 2008)*. Given the importance of sMEK1 in cell function, it is intriguing to discover its potential interaction with IRE1. To further investigate the link between IRE1 and sMEK1, we used coimmunoprecipitation and yeast two-hybrid assays. Gene constructs of IRE1-pcDNA3.1 and sMEK1-GST-pEBG or IRE1-pcDNA3.1 and pEBG-GST were co-transfected into HEK293T cells. Cells were treated with Tunicamycin for 4h to induce ER stress *(Belyy et al., 2020).* Then, immune pulldown was carried out using an anti-IRE1 antibody with lysates from both transfected cells. After immunoprecipitation, the precipitated proteins were immunoblotted using anti-GST. The results from this experiment revealed that sMEK1 interacts with IRE1 (Fig. 1A). Further, to confirm the direct interaction between IRE1 and sMEK1; we carried out a yeast two-hybrid assay. The IRE1 fused to the Gal4 transcription activation domain, and the sMEK1 to the binding domain was introduced into yeast cells. After obtaining independent transformant cells, cultures were spotted on the selection media (His-, Trp-, Leu-). Finally, the transformant cells containing the genes for both sMEK1 and IRE1 grew on selection media, thus proving the direct interaction between the two proteins (Fig. 1B).

**Figure 1.**
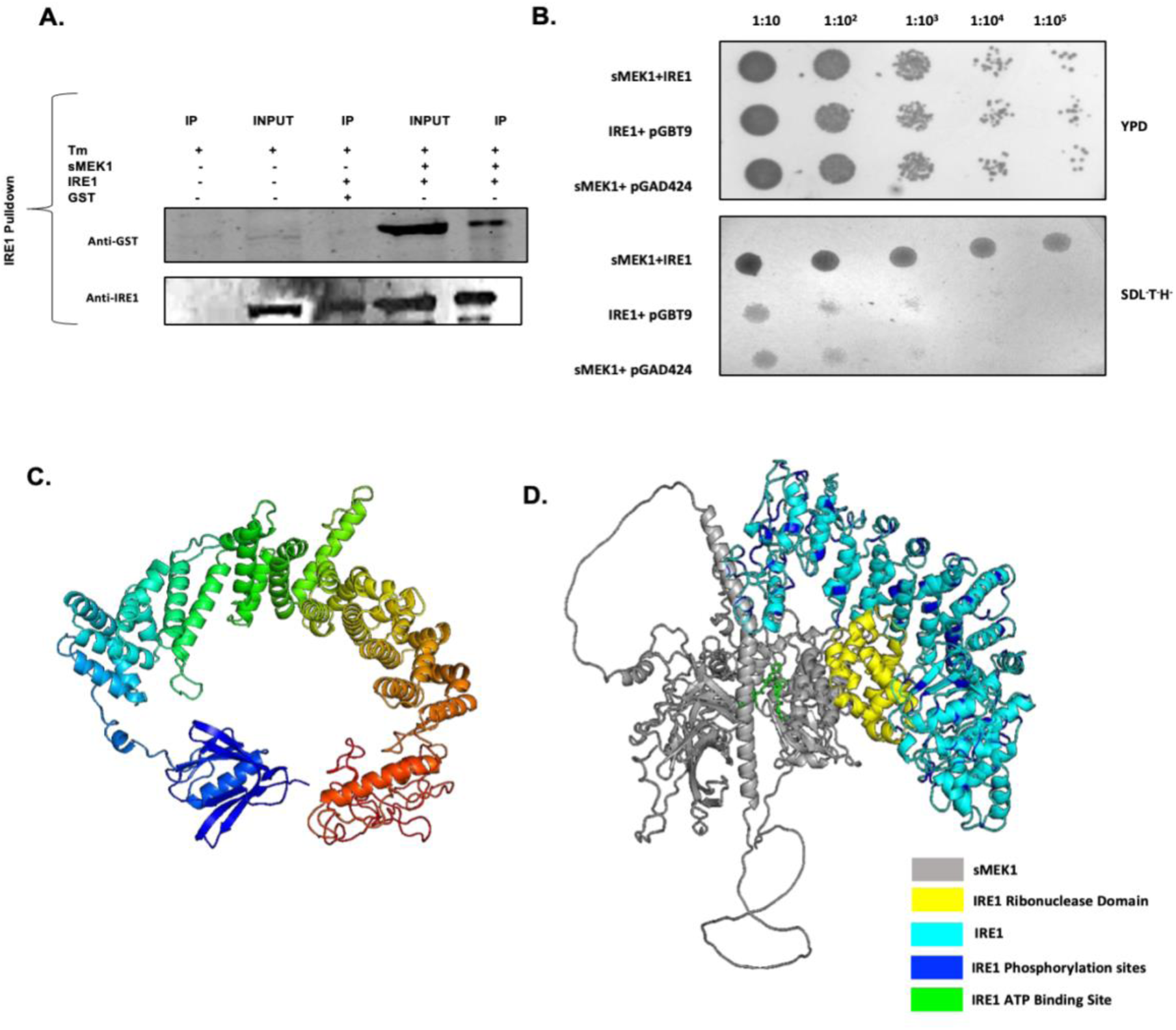
Interaction of sMEK1 with IRE1; **A)** Co-immunoprecipitation of IRE1 with sMEK1. Immunoprecipitation (IP) was performed using an anti-IRE1 antibody and lysates from transfected HEK293T cells. After immunoprecipitation, precipitated proteins were immunoblotted using an anti-GST antibody. lane 1,2 IP and Input for Cells only with no transfection; lane3 IP for GST-pEBG (vector only); lane 4,5 GST-pEBG-sMEK1and pcDNA3.1 IRE1 transfectants. **B)** Yeast Two-Hybrid Assay showing direct interaction of IRE1 with sMEK1. pGAD24-Ire1 + pGBT9-sMEK1 represents colonies obtained from co-transfection of pGAD-IRE1 and pGBT9-plasmids. pGAD24 + pGBT9-sMEK1shows colonies obtained from co-transfection of pGAD24 and pGBT9-sMEK1plasmids. pGAD-IRE1 + pGBT9 represent colonies obtained from co-transfection of pGAD-IRE1 and pGBT9 plasmids. Dilution spotting on SDL^-^T^-^H^-^ drop out media to select for positive interactors (Upper panel). The lower panel shows dilution spotting on YPD rich media. Dilutions were made up to 10^-5^. **C)** Representative picture showing the final model of the sMEK1 protein as deduced from AI-based AlphaFold modelling. **D)** Interaction between sMEK1 annotated in grey model and IRE1. The final docked model is then annotated in terms of phosphorylation sites (Blue), ATP binding sites (Green) and ribonuclease activity in IRE1 (Yellow).

Our interaction studies were further substantiated by computational analysis. To this end, we first recognized the structure of the sMEK1 protein, as no PDB structure of the protein is available to date. We initially used Homology-based approaches to model the sMEK1 protein structure, which couldn’t pose any reasonable template. Thus, Deep learning algorithms like AlphaFold (Supplementary Fig. S1A) and RoseTTAFold (Supplementary Fig. S1B) were employed to model the protein. Among the two models, the former predicted an aligned error structure (Supplementary Fig. S1C). It contained a few loops that were resolved using MD simulation (Supplementary Fig. S1D). After reconciling the low-confidence loops using the DEMO server. We determined the final model for sMEK1, as represented in Fig. 1C. This model was used for further docking analyses with IRE1 (Accession ID-O75460) using the Cluspro server. The final docked model is then annotated in terms of phosphorylation sites, ATP binding sites and ribonuclease activity in IRE1 to see how phosphorylation sites of IRE1 are aligned concerning the interface of the two proteins and how it can affect the docking with a kinase enzyme (Fig 1D). Together these results, including the experimental and bioinformatic analysis, established that sMEK1 directly interacts with IRE1.

### 2.2 IRE1 and Akt are functionally linked by sMEK1 phosphatase

The mammalian target of rapamycin (mTOR) pathway is known to regulate various cellular processes, such as cell growth, metabolism, and autophagy *(Saxton and Sabatini, 2017).* Akt, a serine/threonine kinase, is a central mediator of the mTOR pathway *(Ersahin et al., 2015)*. The interplay between the UPR and mTOR pathways is intricate and likely involves multiple factors and regulatory feedback mechanisms that still need to be fully understood. In this context, we showed how UPR and Akt are functionally linked through IRE1. Our studies validated the connection between IRE1 and Akt pathways through sMEK1 and determined the relay of signal between the three molecules; sMEK1, Akt, and IRE1. We carried out sMEK1 overexpression and knockdown studies in HEK293T cells. Tunicamycin (6μM) for 4h duration induced ER stress. It was found that the treatment with Tm resulted in the dephosphorylation of Akt in the presence of sMEK1(Fig. 2 A), and the phosphorylation restores when cells were knockdown with sMEK1 siRNA (Fig. 2 B), indicating the presence of sMEK1 as a prerequisite to dephosphorylate Akt.

**Figure 2.**
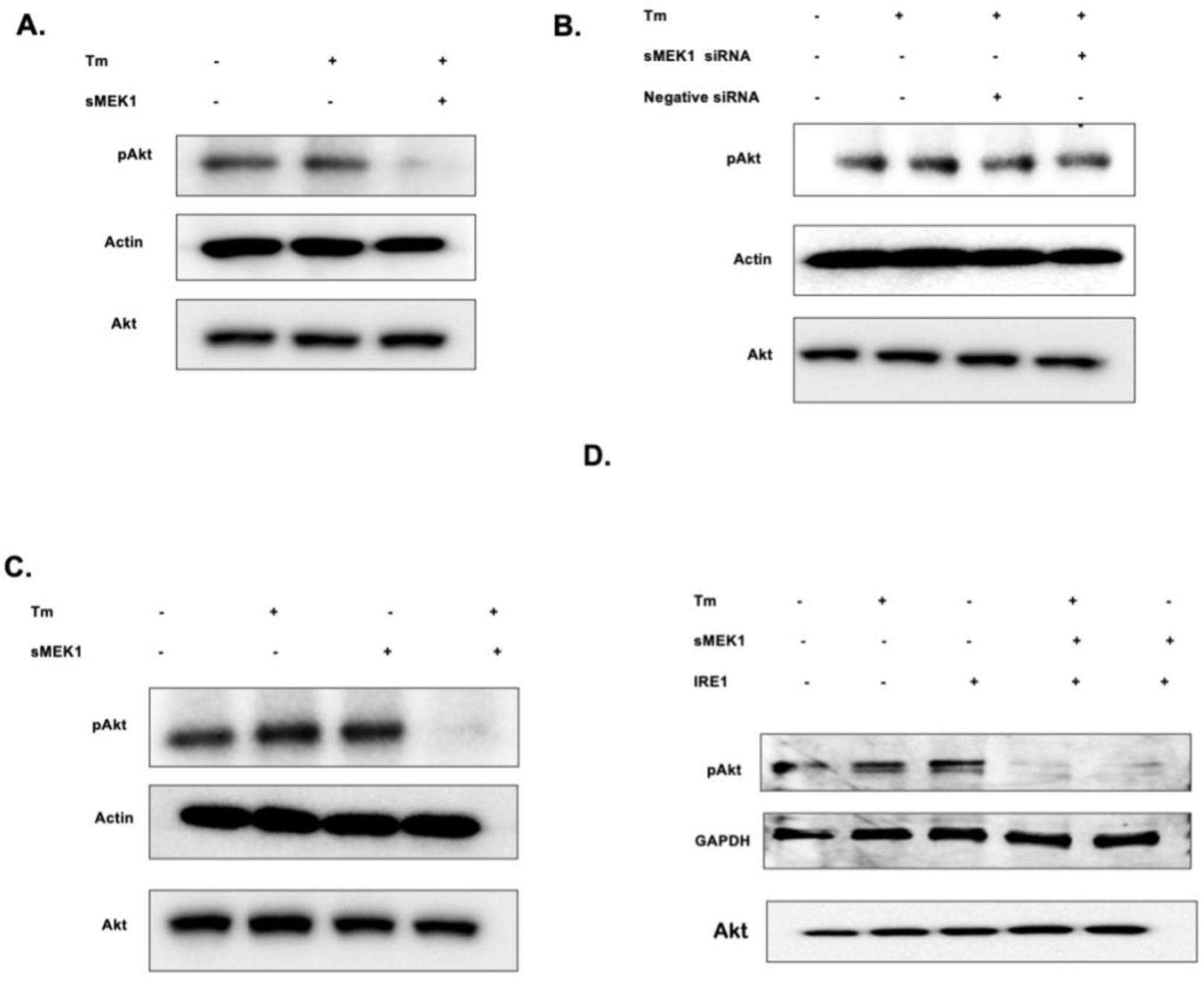
sMEK1 dephosphorylated Akt through IRE1. **A)** Analysis of Akt phosphorylation in sMEK1 overexpressing conditions. HEK293-T cells were transiently transfected for sMEK1-pcDNA3.1. After 36hr cells were stimulated with 6μM tunicamycin for 4hr. Levels of pAkt were checked by probing with Anti-pAkt antibody. **B)** Effect of pAkt under sMEK1 knockdown conditions. Cells were transfected with small interfering RNAs for 24h for effective knockdown of sMEK1, and its effect on pAkt was analysed using western blotting. **C)** Analysis of Akt phosphorylation under different combinations of Tm (6μM, 4h) and sMEK1 overexpression. Akt phosphorylation levels were checked using Anti-pAkt antibody. **D)** Analysis of Akt phosphorylation in IRE1 and sMEK1 overexpressed conditions. Cells were transfected for sMEK1-pcDNA3.1 and IRE1-pcDNA3.1 and Tm (6μM, 4h). Cell lysates obtained from different treatment conditions were used for Immunoblotting against pAkt-antibody. All the Cell lysates subjected to Western Blotting had GAPDH or Actin as an endogenous control.

We also checked for the role of UPR in sMEK1-mediated dephosphorylation of Akt. For this, we transfected cells with sMEK1 in the presence or absence of ER stress (Tm-6μM; 4h) and compared with the untreated cells. The results showed that sMEK1 was only able to modulate Akt phosphorylation when UPR was active as induced by Tunicamycin (Fig. 2C). This confirmed the role of UPR in Akt regulation and highlighted the importance of UPR in the sMEK1-mediated dephosphorylation of Akt.

To further investigate this pathway, we specifically activated the IRE arm of UPR by over-expressing IRE1 (*Han et al., 2009*). We found a decrease in Akt phosphorylation when sMEK1 was expressed in combination with IRE1. Interestingly, this decrease in Akt phosphorylation was seen in presence of IRE1 irrespective of ER stress induction. This indicated that sMEK1 regulation of Akt was through IRE1, and IRE1 overexpression was able to deactivate Akt phosphorylation effectively (Fig. 2D). Overall, these findings suggest that sMEK1 suppresses Akt activity in an ER stress-dependent manner and this regulation is mediated through the IRE1 arm of UPR. These results shed light on the intricate crosstalk between UPR and Akt signalling pathways.

### 2.3. IRE1 Inhibitors dissect the functional role of IRE1 in IRE1/sMEK1/Akt connection

IRE1 protein is peculiar in its dual enzyme activity, kinase and endoribonuclease, and a scaffolding protein that acts as a signalling platform for different binding partners (*Bashir et al., 2021, Han et al., 2009, Urra et al., 2020))*. To ascertain the role of IRE1 in regulating Akt activation, we extended our studies to determine which activity of IRE1 was crucial for the sMEK1 mediated regulation of Akt. To this end, we used IRE1 specific inhibitors that target different functional outputs of IRE1. These IRE1 modulating chemicals are divided into three groups; i) Kinase inhibitors that can differentially activate RNase activity (Type I Inhibitors); ii) Kinase inhibitors that inhibit the RNase activity altogether (Type II Inhibitors/ KIRA, “kinase inhibiting RNase attenuators”); iii) IRE1 RNase domain inhibitors; that directly inhibit the endonuclease activity of IRE1(Xbp1/RIDD) activity *(Ghosh et al., 2014; Lerner et al., 2012; Wang et al., 2012).* We used all three types of inhibitors viz APY29, a type I kinase inhibitor, KIRA6, a type II kinase inhibitor, and STF083010-a small molecular inhibitor of IRE’s RNase.

We conducted a series of experiments to explore the relationship between Akt, and IRE1. Initially, we sought to determine if the kinase activity of IRE1 was vital for sMEK1-induced ATP dephosphorylation. To do so, we used APY29 to inhibit kinase activity and overexpressed sMEK1 in HEK293T cells. We then carried out western blotting for pIRE1 and RT-PCR for RNase outputs. Our findings indicated that APY29 inhibited pIRE1 and divergently inhibited RIDD activity without affecting Xbp1 splicing (Supplementary Fig S2A-C), implying that the kinase activity of IRE1 is shut in a way that its RNase outputs are differentially inhibited (Fig 3A). Under these conditions, we checked how the phosphorylation of Akt is affected using immunoblotting for Akt (S473). Interestingly, it was found that sMEK1 could inhibit p-Akt even in the presence of APY29 (Fig 3A). This suggests that the kinase function of IRE1 is not required for sMEK1 mediated inhibition of p-Akt. Next, we examined whether IRE1’s RNase activity was responsible for the sMEK1-mediated dephosphorylation of Akt. We used RNase inhibitor STF083010 and observed that it effectively inhibited the RNase activity of IRE1 without affecting its kinase domain (Fig 3B, Supplementary Fig S2D-F). We found that just like the kinase activity of IRE1, RNase activity was also not essential for sMEK1-directed Akt dephosphorylation (Fig. 3B). sMEK1 efficiently inhibited Akt phosphorylation even in the presence of STF083010.

**Figure 3.**
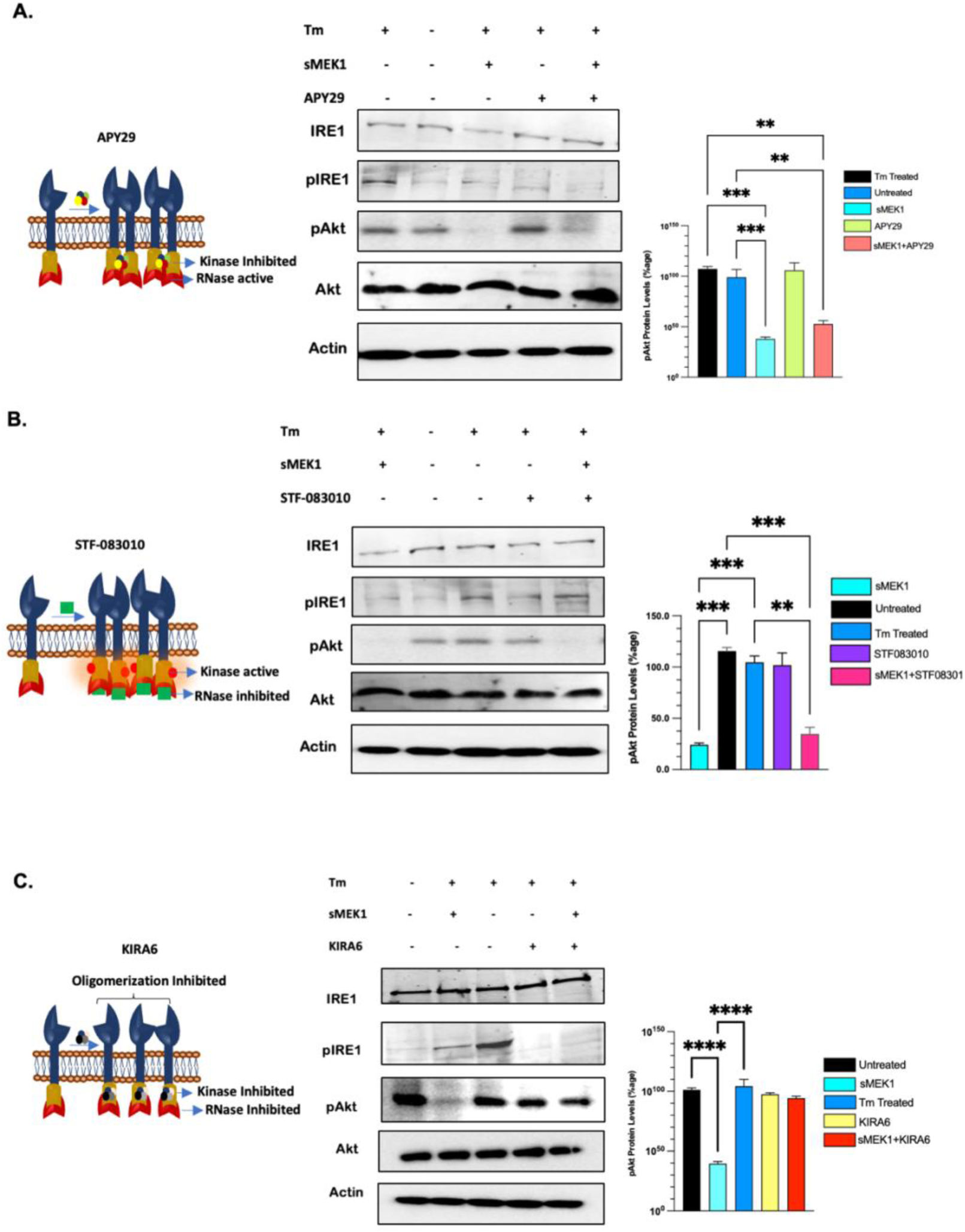
IRE1 Inhibitors dissect the functional role of IRE1 in the IRE1/sMEK1/Akt connection. **A)** Western blot analysis showing the effect of APY29 on pIRE1 and pAkt protein levels compared to their whole proteins and Actin as loading control. **B)** Western blot analysis showing the effect of STF-083010 on pIRE1 and pAkt protein levels compared to their whole proteins and Actin as loading control. **C)** Western blot analysis showing the effect of KIRA6 on pIRE1 and pAkt protein levels compared to their whole proteins. and and Actin as loading control.

Having established that the kinase and RNase activities of IRE1 were not required for inhibition of Akt through sMEK1, we employed KIRA6, a type II kinase inhibitor of IRE1 that destabilizes the molecule such that its kinase, RNase, and oligomerizing activity are inhibited thus rendering the protein non-functional (Supplementary Fig S2G-I). We treated HEK293T cells with KIRA6 and overexpressed sMEK1 under Tm-stimulated conditions. We found that sMEK1 was not able to dephosphorylate Akt in the presence of KIRA6 inhibitor (Fig 3C). This suggests that IRE1 serves as the scaffolding protein necessary for sMEK1 mediated dephosphorylation of Akt. Taken together, our findings indicate that IRE1 is necessary for sMEK1-mediated inhibition of Akt phosphorylation. This effect is due to the scaffold activity of IRE1, ruling out the involvement of its kinase or RNase function.

### 2.4. Akt forms a ternary complex with sMEK1 and IRE1

In continuation with the above observations that indicate sMEK1 interaction with IRE1 resulted in a downstream effect on Akt phosphorylation, implying that the three proteins IRE1, sMEK1 and Akt are in close association. To validate the interaction between IRE1, sMEK1, and AKT proteins, we carried out co-immunoprecipitation and Yeast Two Hybrid Analysis. For Co-IP, gene constructs of IRE1-pcDNA3.1 and Akt-HA-pcDNA or sMEK1-pcDNA 3.1 and Akt-HA-pcDNA were co-transfected into HEK293T cells. Following transfection, 4hr Tm treatment was given to the cells. This was done to ensure that UPR is activated and the interaction of IRE1/Akt or Akt/sMEK1 is captured in its active form. Then, immuno-pulldown was carried out by using anti-IRE and anti-sMEK1 antibodies with lysates from transfected cells. After immunoprecipitation, the precipitated proteins were immunoblotted using an anti-HA antibody. The results from this experiment revealed that there was an interaction between IRE1 and Akt as well as between sMEK1 and Akt, as the band for sMEK1 against IRE1 pulldown was detected (Fig. 4A, C). Furthermore, an inverse IP against HA-Pulldown was carried out that corroborated the interaction between sMEK1 and IRE1 (Fig 4B, D).

**Figure 4.**
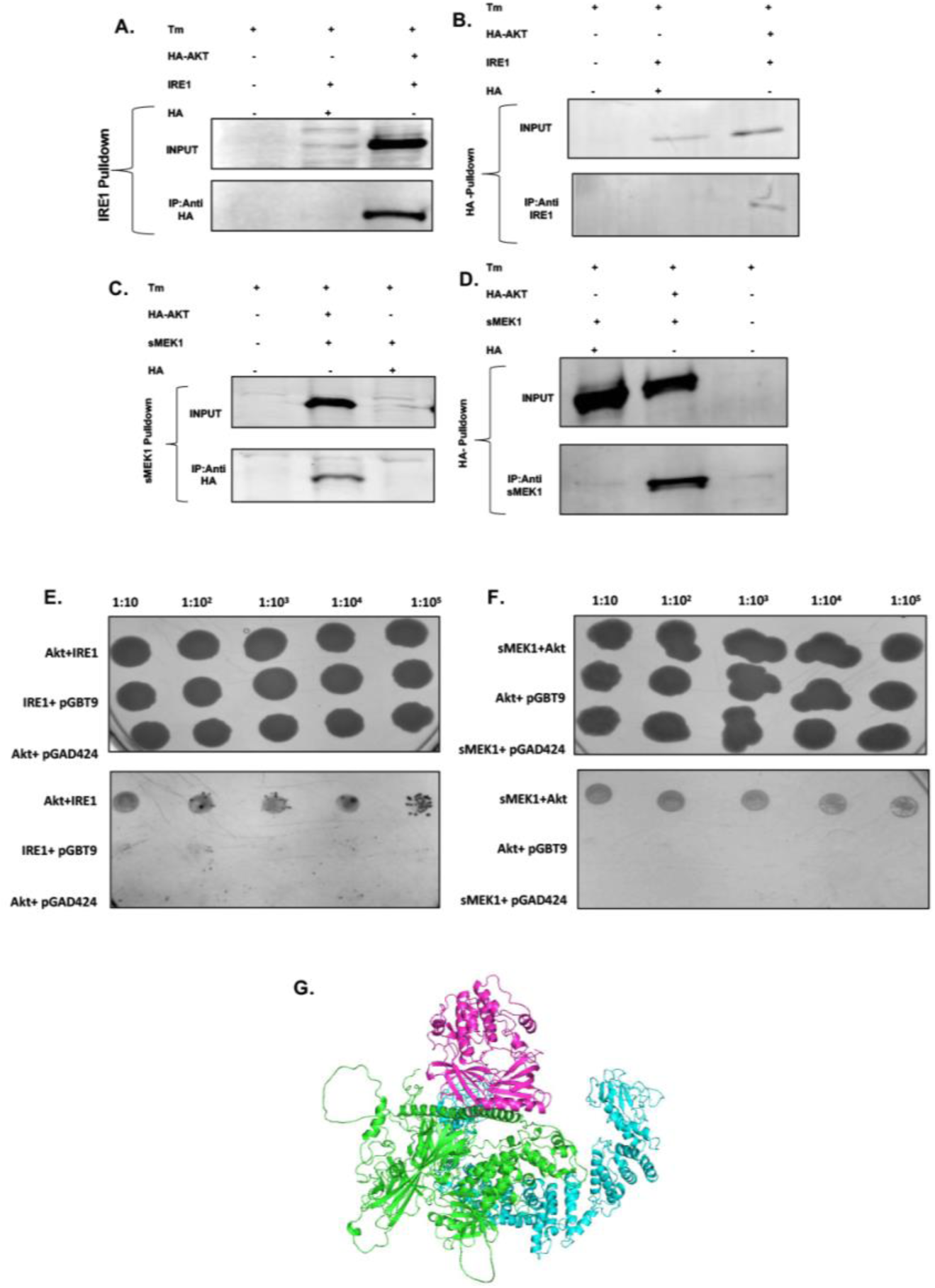
IRE1/sMEK1/Akt forms a ternary complex. A) Co-immunoprecipitation of IRE1 with Akt. Immunoprecipitation (IP) was performed using an anti-IRE1 antibody and lysates from transfected in HEK293T cells. After immunoprecipitation, precipitated proteins were immunoblotted using anti-HA antibody. lane 1, Cells only with no transfection; lane2, HA-pcDNA 3.1 (vector only) ; lane 3 Akt-HA-pcDNA3.1 and pcDNA3.1 IRE1 transfectants B) Inverse IP for IRE1/Akt interaction with HA-pulldown and IRE1 immunoblotting. C) Co-immunoprecipitation of sMEK1 with Akt. Immunoprecipitation (IP) was performed using an anti-sMEK1 antibody and lysates from transfected in HEK293T cells. After immunoprecipitation, precipitated proteins were immunoblotted using anti-HA antibody. lane 1, Cells only with no transfection lane 2 Akt-HA-pcDA3.1 and pcDNA3.1 sMEK1, lane3, HA-pcDNA 3.1 (vector only) transfectants.. Lane1,2 GST-pEBG (vector only)& pcDNA3.1 IRE1, lane3,4 GST-pEBG-sMEK1and pcDNA3.1 IRE1 transfectants, lane 5,6 Input and IP for Cells only with no transfection. D) Inverse IP for sMEK1/Akt interaction with HA-pulldown and immunoblotting against sMEK1 antibody. E, F) Dilution spotting on SDL-T-H-drop out media to select for positive interactors (Upper panel). The lower panel shows dilution spotting on YPD rich media. Dilutions were made up to 10-5. G) Cartoon showing Akt (Purple), sMEK1 (Cyan) and IRE1 (Green) interaction.

**Figure 5.**
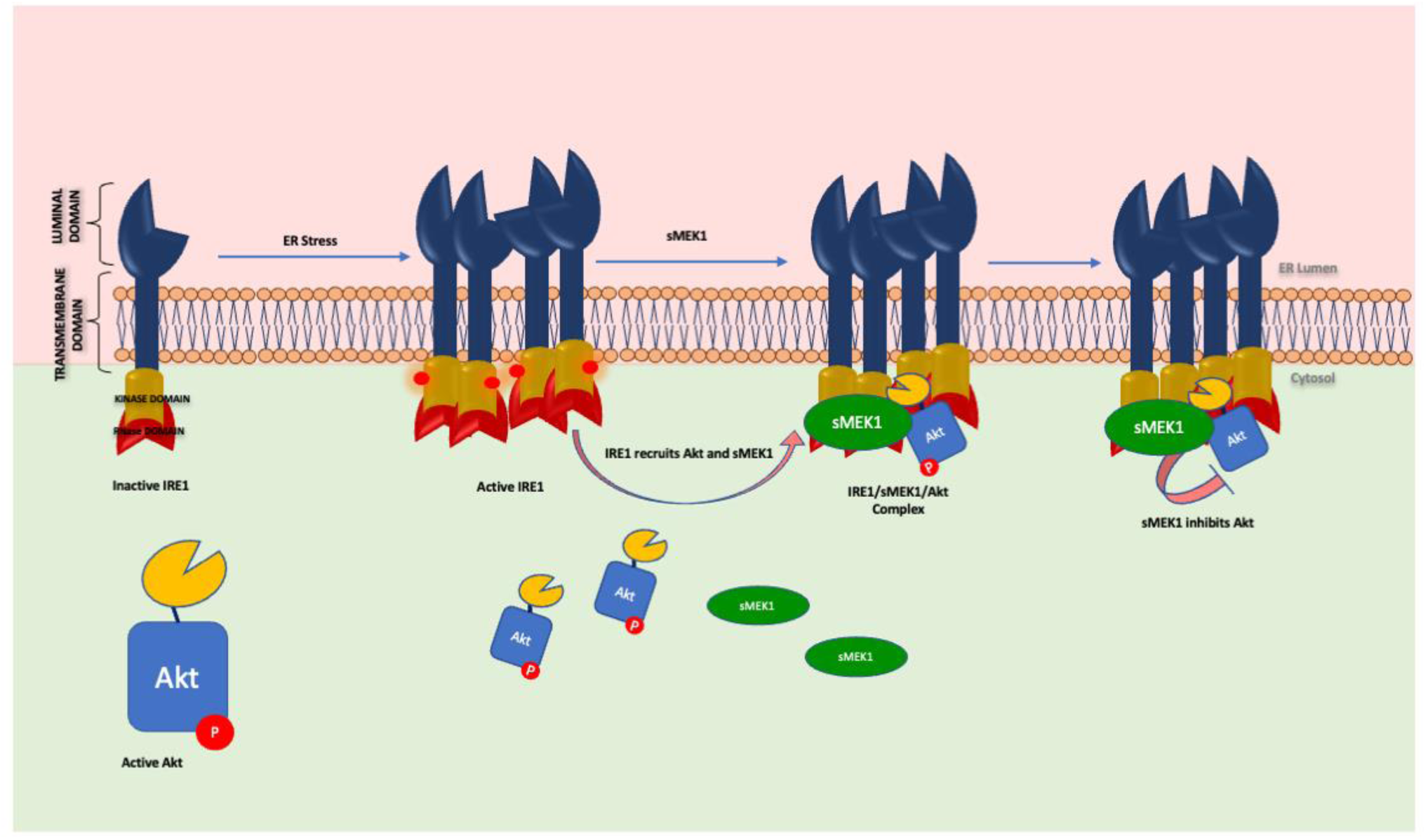
Model for relay of signal from IRE1 sensor of UPR to Akt-signalling via sMEK1. Under ER stress, the monomeric IRE1 comes together to activate its trans-autophosphorylation and, thereby, its endoribonuclease activity. The active IRE1 is able to dock sMEK1 and Akt such that three proteins come in close proximity. This close association of IRE1/sMEK1/Akt leads to the deactivation of Akt by protein phosphatase sMEK1.

To check the nature of interactions between the three proteins IRE1, sMEK1 and Akt, we carried out yeast two-hybrid analyses. To check Akt/IRE1 interaction, AH109 yeast cells were co-transformed with Akt-pGBT9 and IRE1-pGAD424 and selected for co-transformation on SDL^-^ T ^-^ media. After obtaining independent transformant cells, cultures were spotted on the selection media (His^-^, Trp^-^, Leu ^-^). Finally, the transformant cells containing the genes for both Akt and IRE1 grew on selection media, thus, proving the direct interaction between the two proteins (Fig 4E). Likewise, for sMEK1/Akt interaction, sMEK1-pGBT9 and Akt-pGAD424 were co-transformed and selected on SDL-T-media. Cultures were spotted on the His, Trp, and Leu deficient selection media after getting independent transformant cells. Finally, transformant cells carrying both sMEK1 and Akt genes thrived on selection media, demonstrating that the two proteins interact directly (Fig 4F). The results from the above observations imply that the three proteins IRE1, Akt and sMEK1 directly interact with each other forming a ternary complex.

Computational analysis was used to support our interaction studies, whereby we utilized an Alpha Fold model of the sMEK1, IRE1 (Accession ID-O75460), and Akt (Accession ID P31749) structures which had been previously determined. We employed the Cluspro server to conduct further docking analysis and generated a final docked model that was annotated to represent the three proteins. As shown in Fig 4G, the analysis demonstrated that IRE1, sMEK1, and Akt are in close proximity. Taken together with our experimental and bioinformatic analysis, these results suggest that the three proteins may interact with each other in a biological context.

## 3. Discussion

IRE1 acts as a central pathway which orchestrates with different binding partners, leading to the definition of a dynamic signalling platform, ‘UPRosome,’ in which many regulatory and adaptor proteins assemble to activate and modulate downstream responses *(Urra et al., 2020).* These proteins are found to physically interact with IRE1, either serving as its modulators or functional links with other pathways or simply the structural stabilizers of IRE1 *(Hetz and Glimcher, 2009).* Besides the direct physical interactors, there exists functional crosstalk between IRE1 and other cellular signalling pathways *(Urra et al., 2020)*, including one of the important cellular signalling pathways, the mammalian target of rapamycin (mTOR) *(Ozcan et al., 2008; Ozcan et al., 2008; Qin et al., 2010; Sanchez-Alvarez et al., 2017*). Some studies suggest that mTOR selectively inhibits IRE1 activation through AKT-mediated attenuation, influencing IRE1 dynamics, ER-mitochondria contacts, and IRE1 phosphorylation (*Sanchez-Alvarez et al., 2017)*. While, other studies propose that prolonged ER stress negatively modulates mTOR (*Ozcan et al., 2008; Ozcan et al., 2008; Qin et al., 2010*). The idea evolving from these studies is that there exists a functional crosstalk between UPR and Akt pathways which may act antagonistically but integrate and transduce to produce a substantial biological outcome and consequently act in determining the fate of a cell. Despite this, the molecular mediators in this connection remain largely unknown, emphasizing the need for further exploration in unravelling the intricate molecular details.

In this context, one of the proteins revealed by our mass spectroscopy studies is sMEK1 which is found to interact with activated IRE1. Importantly, sMEK1 is known to have a role in regulating the PI3K/Akt/mTOR signalling pathways *(Byun et al., 2012; Kim et al., 2014; Kim et al., 2015)*. sMEK1 is a tumour suppressor protein also known as protein phosphatase 4 regulatory subunit3 (PP4R3), a member of the PP2A subfamily, which has conserved serine/threonine phosphatases *(Chen et al., 2008).* sMEK1 has a role in many important cellular functions, such as apoptotic cell death, microtubule organization, cell cycle arrest, and DNA damage checkpoints (*Chowdhury et al., 2008; Nakada et al., 2008)*. Our previous study revealed that sMEK1 also modulates IRE1 pathway (*Qadri et al., 2024*).

To corroborate our mass spectroscopy data, we validated the interaction between IRE1 and sMEK1 using experimental and computational methods. It turned out that sMEK1 interacts directly with IRE1. Owing to this experimental evidence and cited literature, we sought to analyze the cross-talk between IRE1 and Akt. We started by investigating the effect of ER stress on Akt phosphorylation in the presence of sMEK1. We observed that sMEK1 dephosphorylates Akt only in the presence of Tunicamycin, an ER stress inducer. This suggests that active UPR was important for sMEK1 mediated regulation of Akt. These results support the earlier data that showed sustained ER stress can inhibit the mTOR/Akt pathway, leading to its inactivation and triggering death via autophagy *(Qin et al., 2010)*. Furthermore, our findings indicate that by exclusively activating the IRE1 pathway and not engaging the other three branches of UPR *(Han et al., 2009)*, sMEK1 can inhibit Akt phosphorylation. This suggested that the IRE1 pathway is actually involved in regulating the Akt pathway. It was clear from these experiments that sMEK1 is a key player in driving the cross between IRE1 and Akt. Also, IRE1 is upstream and crucial for sMEK1-mediated regulation of Akt.

Now, the critical question was how IRE1 is involved in this regulation. We used different known inhibitors of IRE1, which suppresses specific IRE1 functions. For example, APY29 inhibits IRE1 kinase activity, and STF083010 inhibits its RNase activity. Using both of these inhibitors, we found that sMEK1 was still able to dephosphorylate Akt, suggesting that neither of these functions of IRE1 is required for Akt regulation. Thereafter, we used KIRA6, an ATP competitive inhibitor of IRE1 that allosterically inhibits kinase and RNase activity and prevents IRE1 from oligomerizing. Interestingly, we found that the KIRA6 resulted in the loss of sMEK1-dependent dephosphorylation of Akt. This showed that the IRE1 scaffold activity is required for Akt regulation. It also suggests that IRE1 acts as a scaffold protein and may be involved in the recruitment of both sMEK1 and Akt. This notion was supported by our immunoprecipitation and yeast two-hybrid assays. We found that IRE1, smEK1, and Akt form a ternary complex that results in the dephosphorylation of Akt.

In conclusion, this study establishes a significant relationship between the IRE1 and Akt pathways has been established, shedding light on the underlying mechanism of this link. The research emphasizes that sMEK1 acts as a molecular bridge connecting these two pathways and that the IRE1/sMEK1/Akt trio functions together. The study clearly demonstrates that ER stress triggers the interaction between sMEK1 and Akt with IRE1, allowing sMEK1 to regulate Akt phosphorylation. These findings provide valuable insights into the regulation, crosstalk, and signal transmission between the two pathways.

By elucidating the molecular mechanism and offering an understanding of the overall impact of this connection, this research underscores the significance of the UPR/Akt link and its potential implications for the pathophysiology of various diseases. This study enhances our understanding of the complex interplay between cellular stress responses and signalling pathways, highlighting potential therapeutic targets for the treatment of various diseases associated with the dysregulation of these pathways. The findings of this study on the IRE1 and Akt pathways could have significant implications for cancer research. The Akt pathway is well-known for its crucial role in cancer cell survival and proliferation, and dysregulation of this pathway is a hallmark of many cancers *(Revathidevi et al., 2019)*. On the other hand, the IRE1 pathway plays a critical role in cellular stress response, including ER stress, which is frequently observed in cancer cells *(Raymundo et al., 2020).* The discovery that sMEK1 regulates Akt phosphorylation by exclusively activating the IRE1 pathway suggests that this pathway could serve as a potential target for cancer therapy. Specifically, targeting the IRE1 pathway using small-molecule inhibitors could be a promising strategy to inhibit Akt activation in cancer cells, potentially leading to decreased cancer cell proliferation and survival.

## 4. Material and Methods

### 4.1 Chemicals and Reagents

HEK (Human Embryonic Kidney)-293T cell line was obtained from the National Centre for Cell Sciences, Pune, India. All other chemicals and reagents, including Dulbecco’s modified Eagles medium (Gibco by Life Technologies, USA), Fetal bovine serum (Gibco by Life Technologies, USA), Penicillin-streptomycin solution (Gibco by Life Technologies, USA), Trypsin-EDTA (Gibco by Life Technologies, USA), Tunicamycin (Tm) (Cal Biochem, USA), APY29 (Cayman Chemicals, USA), KIRA-6 (Cayman Chemicals, USA), STF0830105 (Sigma Aldrich, USA), Kanamycin (Invitrogen, USA), 5-Bromo-4-chloro-3-indolyl phosphate (Amresco, Inc, USA), Nitro-blue tetrazolium (Amresco, Inc, USA), LipofectamineTM 2000 (Invitrogen, USA) were procured from the companies mentioned in the parenthesis. For immunoblotting following antibodies were used; GST (Cell Signaling Technologies, Cat: #2622, 1:5000), IRE1α (Cell Signaling Technologies, Cat: #3294, 1:1000), pAkt (S473) (Cell Signaling Technologies, Cat: #9271, 1:1000), GAPDH (Cell Signaling Technologies, Cat: #2118, 1:1000), HA (Cell Signaling Technologies, Cat: #3724, 1:5000), sMEK1 (Thermofisher, Cat: # A300-840A, 1:500), Akt (Santacruz Biotechnology, Cat: sc-5298, 1:500), pIRE1 (S724) (Novos Biologicals, Cat: NB100-2323, 1:1000), anti-Rabbit IgG Alkaline Phosphatase antibody (Sigma Aldrich, USA), anti-mouse IgG Alkaline Phosphatase antibody (Sigma Aldrich, USA), IRDye Goat anti-Rabbit IgG (LI-CORR Biosciences, USA), IRDye Goat anti-Mouse IgG (LI-CORR Biosciences, USA).

### 4.2 Cell Culture and Treatments

HEK293-T were maintained in Dulbecco’s Modified Eagle’s Medium (DMEM) supplemented with penicillin and streptomycin (each 1% v/v), 10% FBS, grown in the humidified environment of 37°C temperature and 5% CO_2_. ER stress-inducing agent, Tunicamycin (Tm), was reconstituted in dimethyl sulfoxide (DMSO) and added to experimental cell cultures. In all cases, the volume of vehicle control did not exceed 0.02% of the total culture volume. Tunicamycin was stored at a stock concentration of 1mM at -80°C for up to 12 months, and 6μM of working concentration was used to induce ER stress in the cells. IRE1 inhibitors used in the study include; the type I kinase inhibitor APY29, type II kinase inhibitor KIRA6, and the RNase inhibitor STF083010. In this study, 30mM of APY29 stock solution was prepared in DMSO. Cells were treated with 60μM of working concentration for an hour before 4hr Tm treatment *(Wang et al., 2012).* 1mM of stock solution was prepared for KIRA6 in DMSO and used 10μM of working concentration as the least cytotoxic to inhibit IRE1 activity. KIRA6 was pre-treated for 1hr, followed by treatment with Tm for 4hrs *(Ghosh et al., 2014).* Cells were treated with 50μM STF-083010 for 2 hr before adding Tm and allowed to incubate for 4 hrs *(Lerner et al., 2012)*.

### 4.3 Plasmid Constructs

The plasmids used in the study including pEBG-GST, pcDNA 3.1 (+), Akt-HA-pcDNA3.1, HA-pcDNA3.1, pGAD424 and pGBT9 were procured from Adgene. pGEMT-sMEK1 was obtained from SinoBiologicals. For transient gene expression, the full-length human IRE1 and sMEK1 genes were amplified from HEK293T cDNA by PCR and cloned into pcDNA3.1 or pEBG-GST vectors, to make IRE1-pcDNA3.1, sMEK1-pcDNA3.1 and sMEK1-pEBG-GST constructs. For yeast two-hybrid analysis, IRE1 and sMEK1 genes were cloned in pGAD24 and pGBT9 yeast vectors, respectively, while the Akt gene was cloned in both pGAD24 and pGBT9 yeast vectors.

### 4.4 Co-Immunoprecipitation

For co-immunoprecipitation, HEK293-T cells were co-transfected with plasmids, pcDNA3.1 (+) IRE1 and pEBG-sMEK1 or Akt-HA-pcDNA and pcDNA3.1 (+) IRE1 or sMEK1-pcDNA3.1 (+) and Akt-HA-pcDNA; using PEI transfection reagent. Post 36hr transfection, cells were treated with 6μM tunicamycin and incubated for 4 hr. As a control, cells transfected with pcDNA3.1 (+) IRE1 and pEBG empty vector were treated similarly. Cells without transfection were also treated with tunicamycin. Cells were rinsed in phosphate-buffered saline (PBS) and lysed in NETN buffer [50mM Tris-Cl, 150mM NaCl, 1% Glycerol, 1% Np-40, 0.1% SDS, 5mM EDTA, 0.5% Sodium deoxycholate, 10mM NaF, 17.5mM β-glycerophosphate, 1X PIC)]. On the other hand, Protein G plus Agarose beads were washed thrice with NETN buffer and incubated with 2μg of primary antibody (Anti-GST or Anti-IRE1α or Anti-HA or Anti-sMEK1) in 40μL of 1XTBS, overnight on a Nutator (rotating apparatus) at 4°C. On consecutive day, lysates were incubated with anti-IRE1 antibody or anti-GST antibody or HA-Antibody or anti-sMEK1 antibody and precipitated using protein A-agarose (Invitrogen, Carlsbad, CA, USA). The beads solution was washed with NETN buffer and samples were heated with 2X Laemmili Buffer. Precipitated proteins were separated by 10% sodium dodecyl sulfate polyacrylamide gel electrophoresis (SDS-PAGE), and subjected to immunoblot analysis.

### 4.5 Yeast Two-Hybrid Assay

AH109 yeast strain containing *His* gene under GAL4 promoter was used to carry out Yeast Two Hybrid assay. IRE1 and Akt genes were cloned in-frame with the GAL4 DNA binding domain in the vector pGAD24 and, sMEK1 and Akt genes were cloned into the GAL4 DNA activation domain in the vector pGBT9. The AH109 strain was co-transformed with plasmids and selected for Leu and Trp SD-drop out media (SDL^-^T^-^). SDL^-^T^-^Agar was patched with single colonies and cultured at 30°C for three days. Co-transformant colonies were spotted on SDL-T-H (SD complete media with leucine, tryptophan, and histidine dropout) plates to identify protein-protein interactions. Plates were continuously observed for 5-6 days to assess their growth. Apart from the bait clones, empty vectors were transformed as a control for the yeast two-hybrid analysis. Overall, three co-transfections were done for each experiment. Since the promoter of the histidine biosynthesis gene responds to the transcription factor GAL4. As a result, the expression of His-gene will depend on the direct induction of the GAL4 transcription factor via direct interaction between two proteins, allowing colonies to thrive on SDL^-^T^-^H^-^drop out medium.

### 4.6 Gene Silencing by small interfering RNA

To silence target genes, we transfected cells with Sigma Aldrich’s Mission esiRNA using Lipofectamine 2000 reagent per the manufacturer’s instructions. To mitigate non-specific effects, a Mission siRNA fluorescent universal negative control #1 (Cyanine 3) was co-transfected. Cells were grown to 65% confluency in 6cm dishes. The transfection mixture, consisting of 1400ng siRNA in 250μL DMEM (Mixture A) and 5μL Lipofectamine in 245μL DMEM (Mixture B), was prepared, mixed, and added to the cells in serum-free media. After 12-16 hours of incubation, complete media was added, and cells were cultured for 24h and 48h before harvesting. Western blot analysis at 24 hours confirmed target gene knockdown, and all experiments were conducted at this time point.

### 4.7 Western Blotting

Cells were harvested and washed thrice with PBS (pH=7.4), and whole-cell lysates were prepared using the NP40 cell lysis buffer [50mM Tris-Cl, 150mM NaCl, 1% Glycerol, 1% Np-40, 0.1% SDS, 5mM EDTA, 0.5% Sodium deoxycholate] which was supplemented by NaF (10mM), β-Glycerophosphate (17.5mM), and Protease Inhibitor Cocktail (PIC) (1X). Protein concentrations were determined using the Bradford Assay *(Bradford 1976)*. An equal amount of protein (20–30μg) was electrophoresed by 12% SDS PAGE and transferred to a PVDF membrane. The membrane was blocked with 5% BSA in 1XTBS buffer and incubated with primary antibody overnight at 4°C. The blot was then probed against the Mouse or Rabbit Secondary Antibody (*LI-COR Odyssey, USA*) with 1:5000 dilutions. The blot was analyzed using the LI-COR imaging system (*LI-COR Biosciences, USA*). Secondary antibody ALP conjugate (1:40,000 dilution) was also used, and its blots were analyzed on the Gel doc system (*BioRad, CA, USA*).

### 4.8 RNA Extraction and Quantitative PCR

To extract RNA, cells were briefly washed three times with PBS (pH=7.4) and subsequently lysed using TRIZOL reagent. To ensure complete dissociation of nucleoprotein complexes, the samples were allowed to sit at room temperature for 5 minutes. Following this, 200μL of chloroform was added, and vigorous shaking was employed. The samples were then left at room temperature for 15 minutes and subsequently centrifuged at 10,000rpm for 20 minutes at 4°C. The aqueous phase was separated and collected into a different tube containing 500μL of Isopropanol. This resulting mixture underwent another centrifugation at 10,000rpm for 10 minutes at 4°C. The RNA pellet obtained was washed with 70% ethanol and briefly air-dried, after which it was resuspended in an appropriate volume of DEPC-treated water.

For the reverse transcription step, the manufacturer’s instructions for the RevertAid First Strand cDNA Synthesis Kit (Thermo Fisher Scientific, Cat. # K1621) were followed. Subsequently, real-time PCR, performed with 70ng cDNA and the SYBR Green PCR Master Mix (Applied Biosystems, Foster City, CA, USA), was carried out using duplicate reactions for each sample in a 96-well plate. To account for variations, gene expression was normalized to the housekeeping gene β-Actin as an endogenous control. The following primer sets were used in the study are mentioned in Table 1. These reactions were conducted in the 7500 Real-Time PCR System from Applied Biosystems, and quantitation of each transcript was achieved by normalizing the comparative threshold cycle with endogenous controls.

**Table 1.**
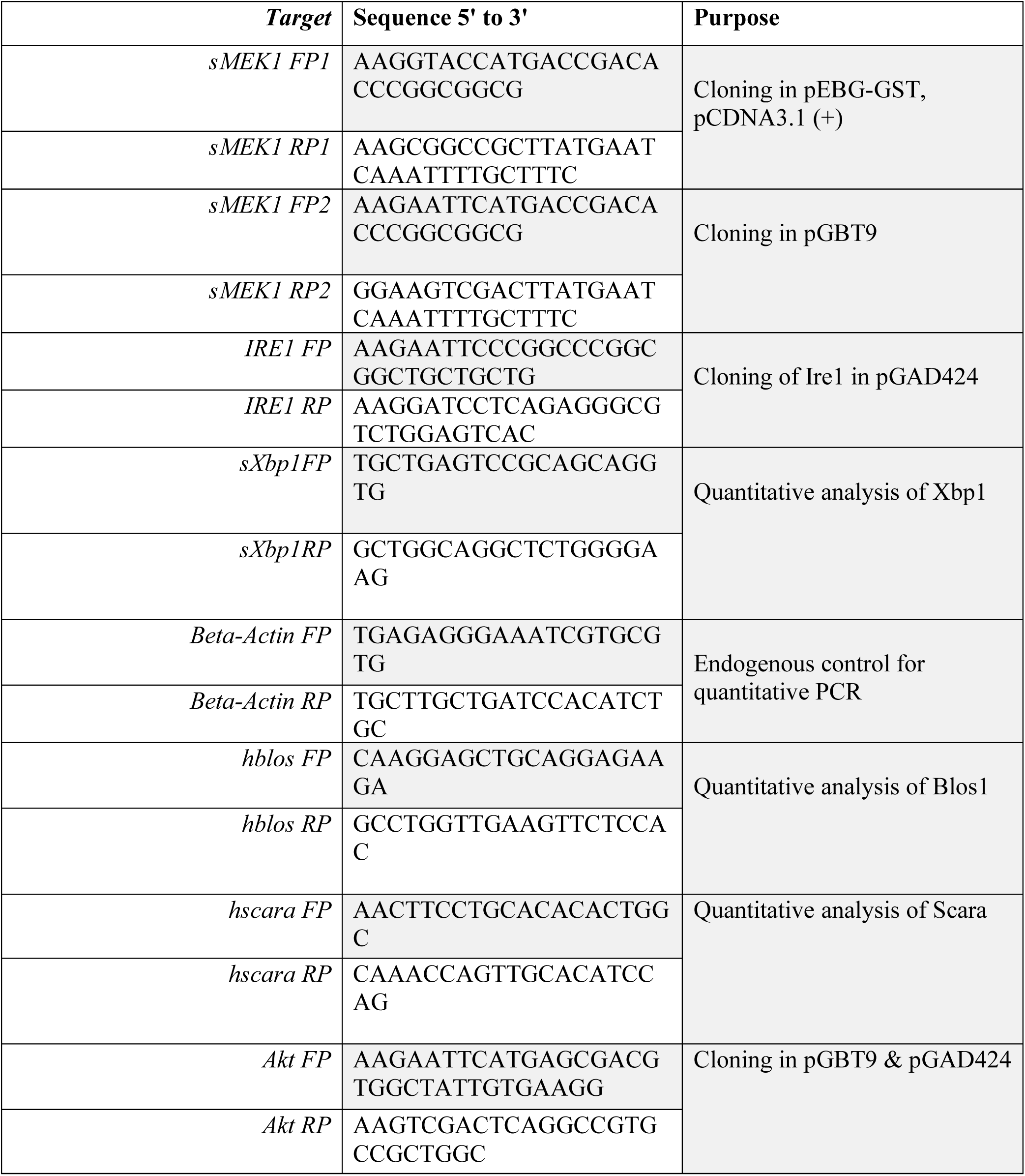
Primers used in the study.

### 4.9 Computational Protein Modelling

Homology modelling and Deep learning approach were used to determine the sMEK1 structure. For the Homology based modelling of sMEK1, DomSerf *(Buchan et al., 2013)* was used. DomSerf is a fully automated, protein-domain homology modelling pipeline combining pDomTHREADER, PSI-BLAST, DomainFinder and MODELLER. Alpha Fold *(Jumper et al., 2021)* and RoseTTA Fold *(Baek et al., 2021)* were used in the deep learning approaches. Alpha Fold is an AI system developed by DeepMind that predicts a protein’s 3D structure from its amino acid sequence. In contrast, RoseTTA Fold accurately predicts protein structures and interactions using AI deeplearning models and a three-track neural network.

### 4.10 ClusPro Server Protein-Protein Docking

ClusPro is the first completely automated, web-based software for docking protein structures computationally. It requires either utilizing ClusPro’s web interface to upload the coordinate files of two protein structures or inputting the relevant PDB codes, which ClusPro will then download from the PDB server. Before picking a limited number of complexes with acceptable surface complementarities, the docking algorithms assess billions of candidate complexes. The set of structures is then screened for further clustering using a process that picks those with good electrostatic and desolvation-free energies. The program provides a selection of putative complexes that are graded and automatically supplied to the user based on their clustering properties.

### 4.11 Statistical analysis

The results of this work are reported as means of ± SEM. The significance of differences between groups was determined using an unpaired t-test and one-way ANOVA. p< 0.05 was considered statistically significant. The statistical analysis was performed using GraphPad PRISM v6.03 statistical software (*GraphPad Software, La Jolla, CA*). Densitometry analysis was done using ImageJ software *(NIH)*.

## Author contributions

Conceptualization: OQ, SB and KMF; Methodology: OQ, SB, MB and DP; Formal analysis and investigation: OQ; Writing – original draft preparation: OQ and KMF; Writing– review and editing: OQ, SB and MB; Funding acquisition: KMF; and Supervision: KMF

## Acknowledgement

We are thankful to Indian Council of Medical Research (ICMR), Department of Science and Technology, Ministry of Ayurveda, Yoga & Naturopathy, Unani, Siddha and Homoeopathy (AYUSH), Government of India, and, for providing individual fellowships. We extend our gratitude to the Department of Science and Technology (DST_SERB), New Delhi, India for lab funding.

## Notes

### Competing Interest Statement

The authors have declared no competing interest.

### Summary of Updates

We have thoroughly revised the paper, incorporating several significant updates. The author list has been updated to include a new contributor. The abstract has been revised to reflect the new content and findings of the paper. Additionally, the Introduction has been aligned with the updated scope of the study. The Results sections have been renamed for clarity. We have added additional data to Figure 1, updated the Western blot images in Figure 2, and included more data in Figure 3. A new Results section has been introduced, with the previous Results 3 reorganized as Results 4 and the new data presented as Results 3. The figure illustrating the model has also been updated to accurately represent the revised findings. The Discussion section has been extensively revised to incorporate and contextualize the new data. The Materials and Methods section has been expanded and detailed to provide a clearer description of the methodologies used. The references have been updated to include recent relevant literature. A separate Supplemental file has been added to support the main text, and

