## Supplemental File for "sMEK1 promotes crosstalk between IRE1 and Akt signalling pathways: Evidence for a novel IRE1/sMEK1/Akt complex"

### Supplementary Data

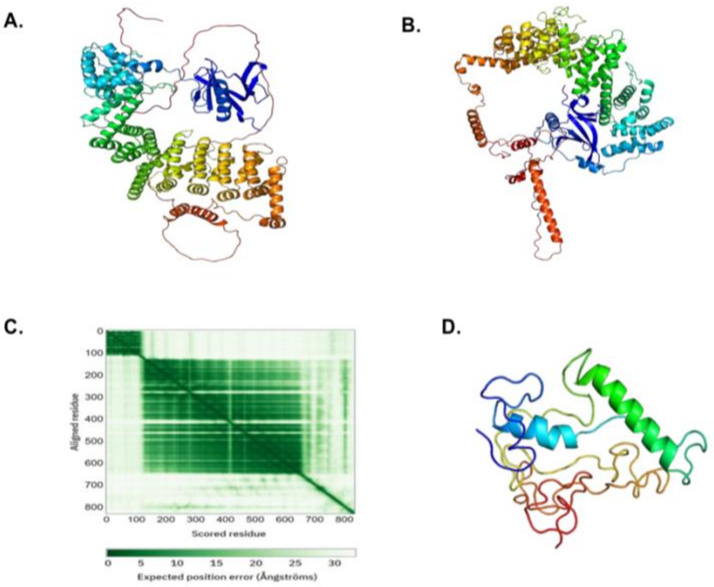


**Figure S1** A) Cartoon diagram showing the predicted structure of sMEK1 using deep learning-based approach Alpha Fold. B) Cartoon showing the predicted structure of sMEK1 using deep learning-based approach RoseTTA Fold. C) Predicted aligned error associated with the AlphaFold predicted model of sMEK1. D. A frame of the MD simulation of the 654-833aa region of sMEK1.

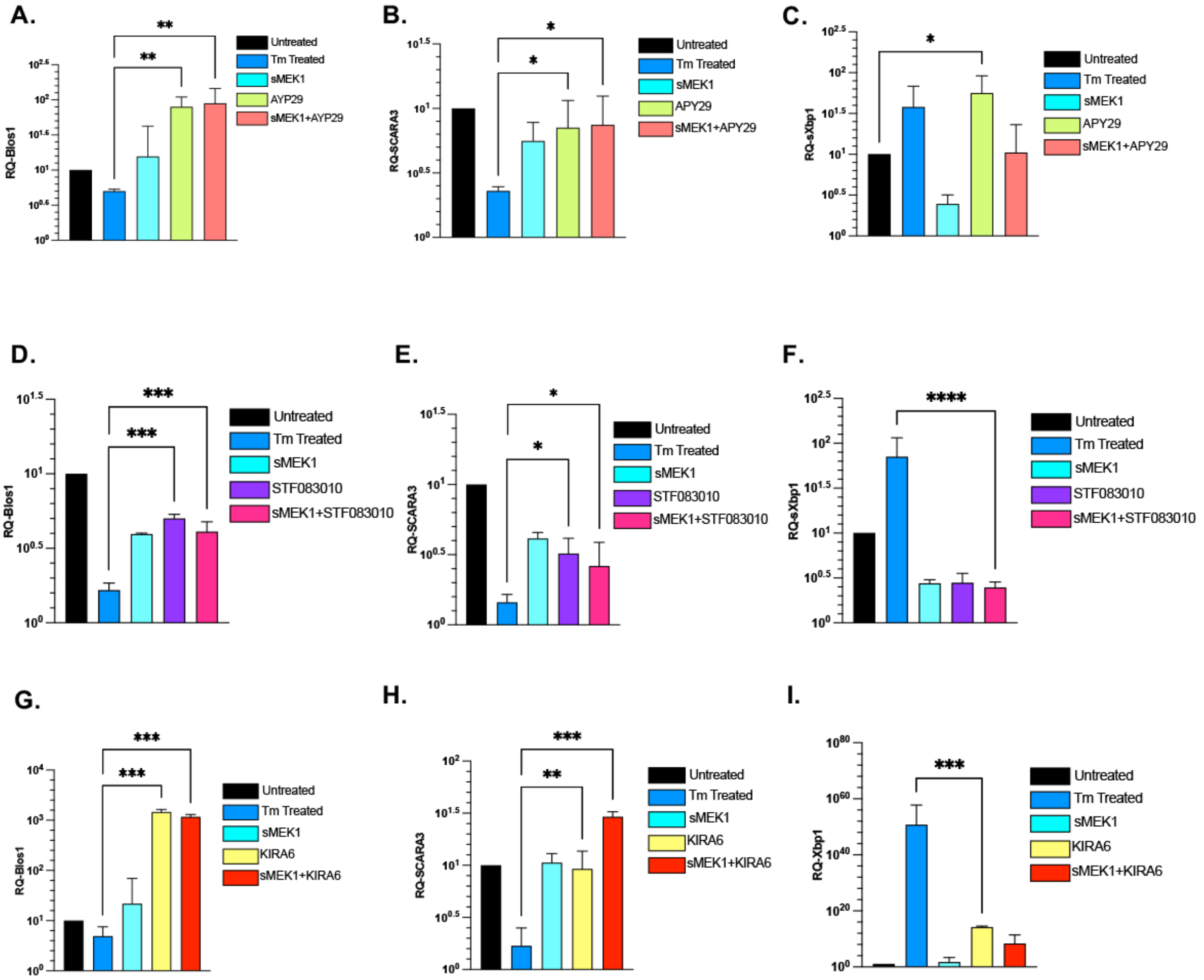


**Figure S2.** A-C) RT-PCR showing the effect of APY29 on the downstream effectors of IRE1; RIDD as shown by the levels of Blos1 and SCARA mRNA; and Xbp1splicing as shown by the levels of sXBP1 mRNA (n=2). (*- $p < 0.05$, **- $p < 0.01$). D-F) RT-PCR showing the effect of STF-083010 on the downstream effectors of IRE1; RIDD as shown by the levels of Blos1 and SCARA mRNA; and Xbp1splicing as shown by the levels of sXBP1 mRNA (n=2). (*- $p < 0.05$, ***-$p<0.001$, ****-$p<0.0001$). H-I) RT-PCR showing the effect of KIRA6 on the downstream effectors of IRE1; RIDD as shown by the levels of Blos1 and SCARA mRNA; and Xbp1splicing as shown by the levels of sXBP1 mRNA (n=2). (**- $p < 0.01$, ***-$p<0.001$).

**Table S1** Mass Spectroscopic data showing IRE1 interacting partners.

| Gene symbol | Gene ID | Description | Coverage | MW [kDa] | calc. pI | Score Mascot |
|---|---|---|---|---|---|---|
| HSPA4L | 22824 | heat shock 70 kDa protein 4L isoform 2 | 5.057471 | 97.597 | 5.95 | 106.0278 |
| EIF3B | 8662 | eukaryotic translation initiation factor 3 subunit B | 9.336609 | 92.424 | 5 | 359.231 |
| HSPH1 | 10808 | heat shock protein 105 kDa isoform 3 | 4.186047 | 97.398 | 5.72 | 112.9489 |
| SRP72 | 6731 | signal recognition particle subunit SRP72 isoform 1 | 6.85544 | 74.56 | 9.26 | 379.3827 |
| SMEK1 | 55671 | serine/threonine-protein phosphatase 4 regulatory subunit 3A isoform X1 | 6.3625450180 | 95.308 | 4.94 | 40.34 |
| RPS8 | 6202 | 40S ribosomal protein S8 | 15.86538 | 24.19 | 10.32 | 247.8473 |
| GCN1 | 10985 | eIF-2-alpha kinase activator GCN1 | 1.198053 | 292.524 | 7.43 | 255.3575 |
| RPS4X | 6191 | 40S ribosomal protein S4, X isoform X isoform | 9.885932 | 29.579 | 10.15 | 98.85069 |
| LETM1 | 3954 | LETM1 and EF-hand domain-containing protein 1, mitochondrial precursor | 2.70636 | 83.302 | 6.7 | 371.1954 |
| RBM25 | 58517 | RNA-binding protein 25 | 3.795967 | 100.124 | 6.32 | 55.34 |
| HNRNPA3 | 220988 | heterogeneous nuclear ribonucleoprotein A3 | 5.820106 | 39.571 | 9.01 | 150.6721 |
| ZFR | 51663 | zinc finger RNA-binding protein | 2.607076 | 116.939 | 9.04 | 39.65 |
| CSDE1 | 7812 | cold shock domain-containing protein E1 isoform 4 | 2.014218 | 93.683 | 6.52 | 22.47 |
| HEATR1 | 55127 | HEAT repeat-containing protein 1 | 0.466418 | 242.215 | 6.54 | 38.85 |
| MRPS5 | 64969 | 28S ribosomal protein S5, mitochondrial | 2.55814 | 47.976 | 9.92 | 43.73 |
| SEC63 | 11231 | translocation protein SEC63 homolog | 1.052632 | 87.942 | 5.31 | 36.82774 |
| EIF4B | 1975 | eukaryotic translation initiation factor 4B isoform 1 | 2.11039 | 69.657 | 5.67 | 81.75 |
| SLC25A24 | 29957 | calcium-binding mitochondrial carrier protein SCaMC-1 isoform 1 | 1.677149 | 53.32 | 6.33 | 41.58 |
| RBM28 | 55131 | RNA-binding protein 28 isoform 1 | 1.317523 | 85.685 | 9.22 | 34.39 |
| HSPA14 | 51182 | heat shock 70 kDa protein 14 isoform 1 | 2.1611 | 54.76 | 5.59 | 28.41 |
| CDC37 | 11140 | hsp90 co-chaperone Cdc37 | 2.380952 | 44.44 | 5.25 | 24.86 |
| LBR | 3930 | lamin-B receptor | 1.300813 | 70.658 | 9.36 | 45.86 |
